# Operational tolerance is early acquired and long maintained in 50% liver allograft transplantation

**DOI:** 10.1101/2022.02.10.480000

**Authors:** Guoyong Chen, Gaofeng Tang, Huibo Zhao, Sidong Wei, Xiaoyan Guo, Fangzhou Liu, Di Lu, Hui Guo, Shaotang Zhou

**Affiliations:** 6th department of hepatopancreaticobiliary Surgery of Henan Provincial People’s Hospital, People’s Hospital of Zhengzhou University, Zhengzhou, 450003, China; Institute of Organ Transplantation, Tongji Hospital, Tongji Medical College, Huazhong University of Science and Technology, Wuhan, 430030, China, Key Laboratory of Organ Transplantation, Ministry of Education, NHC Key Laboratory of Organ Transplantation, Key Laboratory of Organ Transplantation, Chinese Academy of Medical Sciences; Animal center of Henan Academy of Traditional Chinese Medicine, Zhengzhou, 450003, China; Gastroenterology department of Henan Provincial People’s Hospital, People’s Hospital of Zhengzhou University, Zhengzhou, 450003, China

## Abstract

Lifelong anti-rejection therapy is mandatory to weaken the host immune system and maintain the graft functions after organ transplantation, its toxicities and side-effects incidentally elicit mortalities and morbidities. Operational tolerance is a long immunosuppression-free state in which the allograft functions well and no immunological rejections occur, and no operational tolerance is clinically in use. Here we introduce that operational tolerance, based on hypertrophy to hyperplasia switch upon liver regeneration, is early achieved and maintained well in half-size liver allograft transplantation through host bone marrow stem cells repopulation and 9-day immunosuppression. Compared with whole and partial rat liver transplantations as the controls, longer-term survivals (over 430 days) were observed in the tolerant hosts (*p*= 0.001), the allografts functioned well and no acute or chronic rejections were confirmed by histology examinations. Further study revealed that the allograft was reinstituted by host-derived stem cells marked with CD34, which migrated, repopulated and differentiated after mobilization. However donor-specific hyporesponse was not achieved through skin transplantation, indicating no adaptive immune activity occurrence. Our protocol is characteristic of targeting the allografts and almost no immunological intervention, offering a novel avenue to operational tolerance induction which is of highly clinical relevance.

## Introduction

Liver transplantation (LT) is well accepted as the only life-saving treatment for benign end-stage liver diseases, metabolic liver diseases and liver tumors. Chronic immunosuppression is standard-of-care to preserve the donor allografts, bringing about the disreputably adverse effects associated with systemic toxicities, lower quality of life and greater cost burden^1–3^. It was reported in the massive data that the recipients had hardly gained longer-term survival post-LT for decades^2^. Operational tolerance can overcome major histocompatibility complex (MHC) barriers and permit complete withdrawal of immunosuppressants; it holds a promise as the Holy Grail. Peter Medawar first realized skin graft tolerance using fetal mice with immature immunology^4^. Calne et al reported spontaneous tolerance induction using the liver allograft heterotopically placed in pigs^5^. Later Houssin et al also successfully induced tolerance using an accessory semi-allograft in rats while the host total liver removal was performed after the graft implantation to spontaneously accept other grafts like the heart and skin from the same donor, demonstrating that the liver is immunologically privileged^6^. Both protocols did not closely mimic clinical practice. Great immunology progresses have advanced our understanding of rejection as a bi-directional immune activity in nature between the host and the graft. All strategies of tolerance induction targeted the host and immune reaction, deletion or anergy of T cells, blockade of costimulatory signals and adoptive transfer of regulatory T cells were designed and carried out^7,8^,, these proof-of-principle studies cannot be extrapolated to the clinic. Starzl pioneered human LT and pinpointed out chimerism was the frequent state of the long-living recipients^9^. Currently it is documented that macrochimerism achieved by hematopoietic cell transplantation from the solid organ donor is essential to operational tolerance, but maintaining macrochimerism is challenging due to continuous immune deletion or destruction by the host over time, and hematopoietic cell transplantation will incur life-threatening complications like severe infections and graft versus host disease^10,11^, it is noted that clinical trials for this protocol are under way.

Feng et al reported a higher rate of spontaneous tolerance of parental partial allograft in highly selective pediatric recipients, suggesting that liver regeneration was possibly attributable to tolerance^12^. The liver has a unique regenerative capacity to restore his native mass following major resection, this unique phenomenon like an elephant in the room is neglected in tolerance induction. Liver regeneration exists in three forms or sources: hepatocytes (accounting for 80% of parenchyma cells), liver-resident progenitors (<2%) and extra-hepatic stem cells. Generally, the basic premise of hepatocyte proliferation is the intact hepatocytes upon hepatectomy; when intra-hepatic cell proliferation was refrained, extra-hepatic progenitors contributed to this process^13–15^. Here we deployed unique liver regeneration to induce operational tolerance in the setting of half-size liver allogafts in rats, leaving the host and its immune system almost intact.

## Experimental design and grouping

Teleologically our study is to establish the protocol for operational tolerance induction and make sure of its potentially clinical application. Reduced-size (50,70%) allograft LT was frequently performed clinically and used to trigger liver regeneration in our study. 30% liver graft is highly responsible for small-size syndrome and abolished in our study (see Methods). According to the references and our pilot study^16^, whole and 50% graft LT were performed to confirm the effect of liver regeneration of the smaller graft with once-a-day subcutaneous injection of cyclosporine A and recombinant human granulocyte stimulating factor(r-GSF)(see methods)^17^; 50% graft LT was performed as the control only with the same immunosuppressant use to determine the effect of bone marrow stem cells (Table 1, Extended Data Table 1, Extended Data Fig. 1). The experiments are conducted in compliance with the standards and rules for animals set by the Institutional Animal Care Committee of Henan Provincial People’s Hospital.

**Table1.** Experimental grouping for graft size, treatment and survival. Cyclosporine A: CSA; recombinant human granulocyte stimulating factor: r-GSF;* for the death unrelated to rejection. For control groups, the hosts living over 2 weeks were included for survival analysis, the rats living more than 2 months were included for the tolerant group.

| Combinations | Graft | N | Treatment | Survival (days) |
| --- | --- | --- | --- | --- |
| Lewis→BN (ctrl- group 1) | 100% | 5 | CSA+ r-GSF | 123,72,61,65,43 |
| Lewis→BN (ctrl- group 2) | 50% | 6 | CSA | 163,97,157,89,68,59 |
| Lewis→BN(tolerant group) | 50% | 12 | CSA+ r-GSF | 153,167,189,243,237,365,341*,113*,430,263,235,239 |

### Survivals, functions and structures of the rat liver allografts

Clinically operational tolerance is defined as survival of an allograft without chronic rejection in the absence of any immunosuppression for more than 1 year (2 months for small animals). For the nonhuman organ transplantation, the graft survival is equivalent to the host life. In this study graft survival time was calculated from the day of transplantation to the time that chronic rejection was confirmed or the date of death correlated with rejection according to Banff criteria^18^. The core of operational tolerance is to assess whether the allograft’s functions and structures are maintained normal after complete immunosuppression withdrawal. Liver allograft survival, functions and histological examinations were primary end-point foci. Glutamic-pyruvic transaminase (ALT) and glutamic-oxaloacetic transaminase (AST) are conventional parameters to evaluate liver functions. Our result showed that the tolerant rats were freely living much longer than other rats in controls (*p*=0.001, Fig.1). In 14 days post-LT, ALT and AST were gradually becoming normal in all groups; afterwards two values were maintained normal in the tolerant rats, much lower than those of 2 controls along with survival (Fig.2). Histology revealed that acute or chronic rejection developed within 2 months for the controls, and that the normal structures of liver grafts only with mild ductular reaction (scored 1) were detected in the tolerant rats, furthermore; sequential HE staining showed no fibrosis and no acute or chronic rejection in the tolerant rats, which were detected in controlled rats due to rejection (Fig.3,4, Extended Data Fig.2). Immunoglubin G was detected positive in the control groups (Fig.4, Extended Data Fig.2), CK19 as a biomarker of bile duct proliferation was stained more positive in the control rats than that in the tolerant rats (Fig.5, Extended Data Fig. 2).

**Fig.1.**
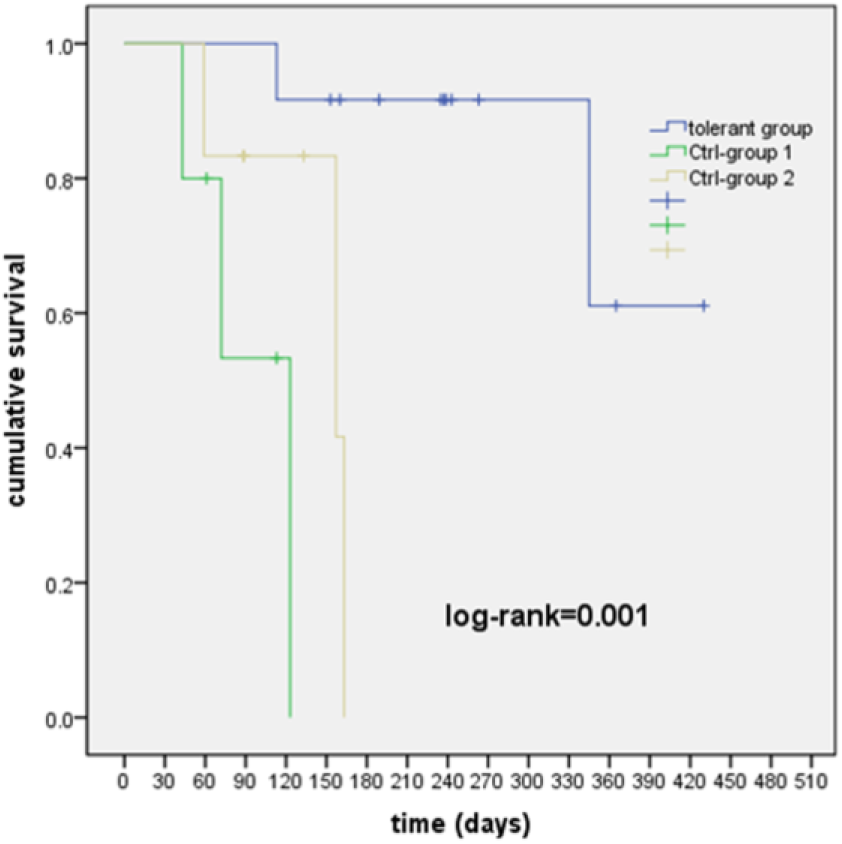
Survival curves for 3 groups evaluated by Kaplan-Meier analysis. The deadline of survival time of all rats for this study was Dec 31.2021.

**Fig.2.**
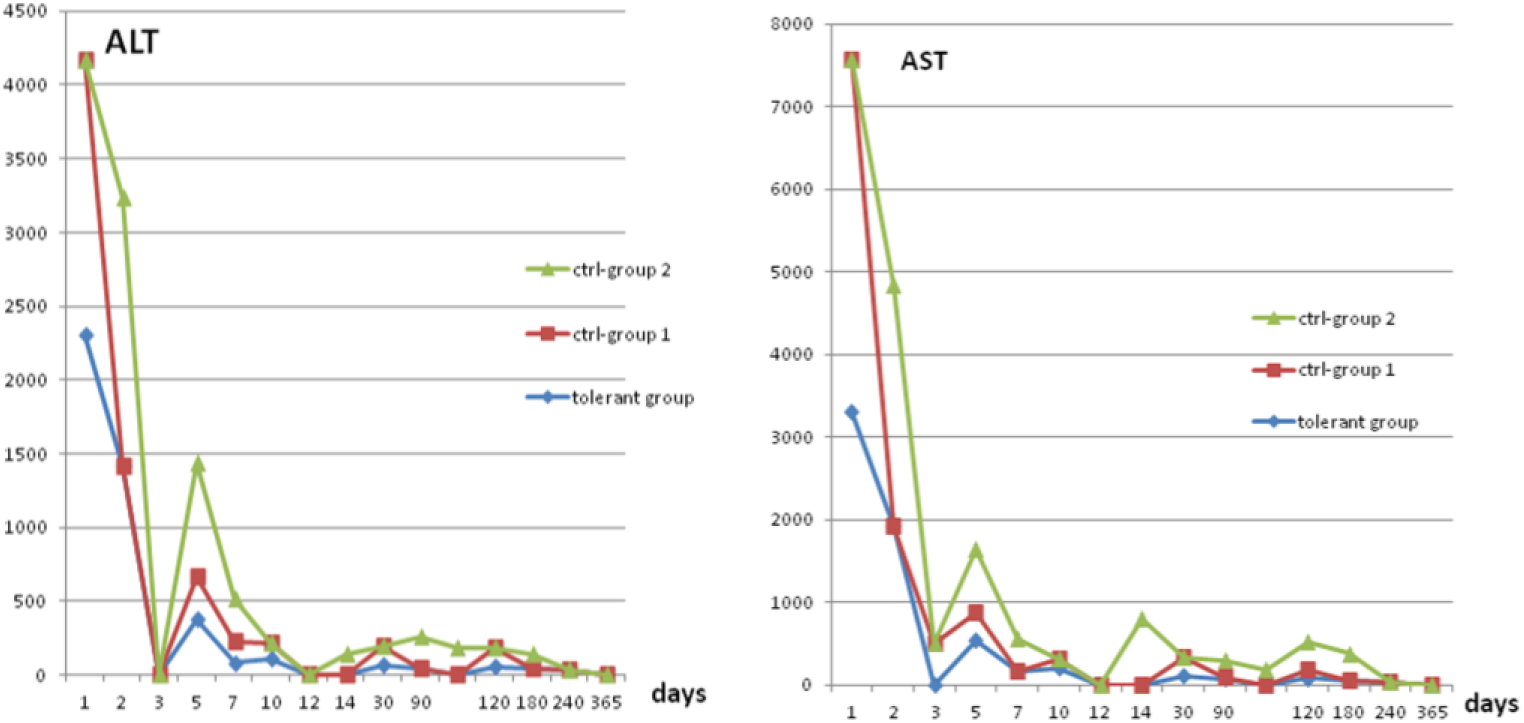
Trends for allograft liver functions with Stacked line chart. ALT, AST for 3 groups, both peak in the early days, were gradually normal in about two weeks, afterward they were going down but higher in the control rats, almost normal all the time only in the tolerant rats,.

**Fig.3.**
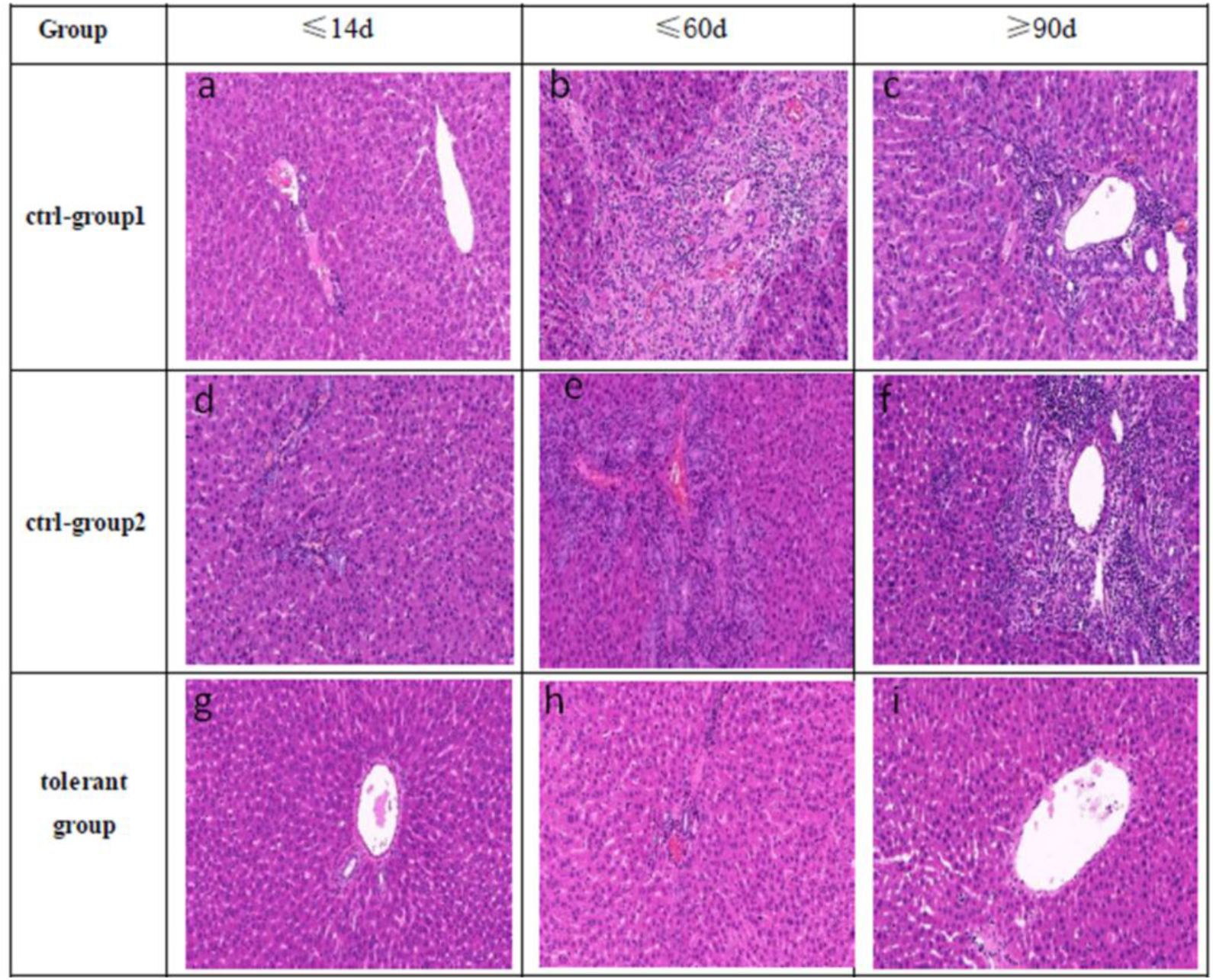
Histology examinations of groups for different time. The grafts were procured at end-point time (death or chronic rejection),a-c, whole graft control group (7,43,123 days), d-f, half size graft control group (6,59,157days). g-i, tolerant group (7d, 53d, 365days)

**Fig.4.**
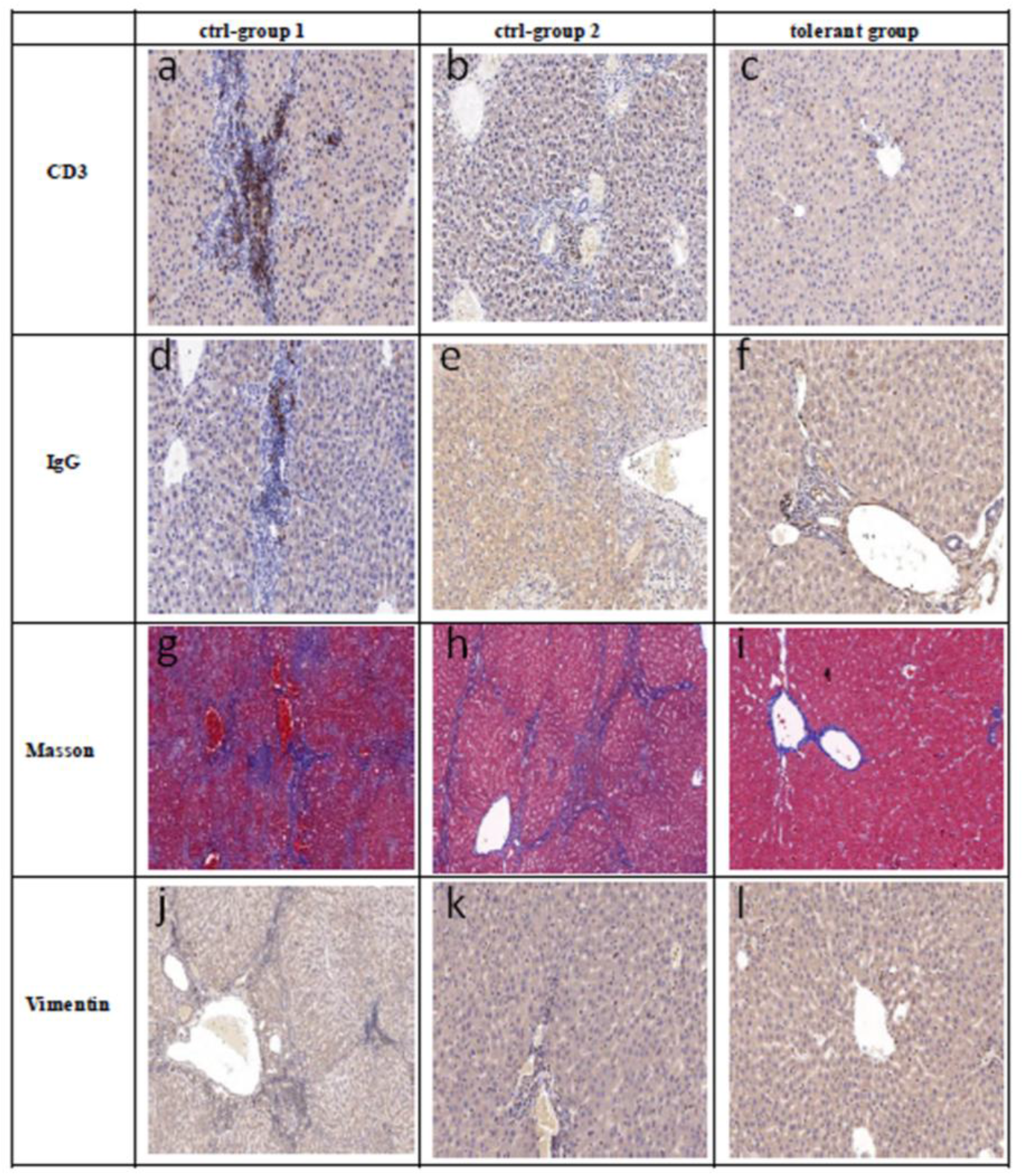
Histoimmunochemical staining for the structures of allograft of 3 groups. CD3 for infiltrated lymphocytes, a (123d),b (157d), c (365d); IgG for immunoglobulin G deposit, d (65d),e (61d),f (341d); Masson, g (59d), h (68d), I (59d) and Vimentin staining for fibrosis, j (59d), k (157d), l (365d).

**Fig.5.**
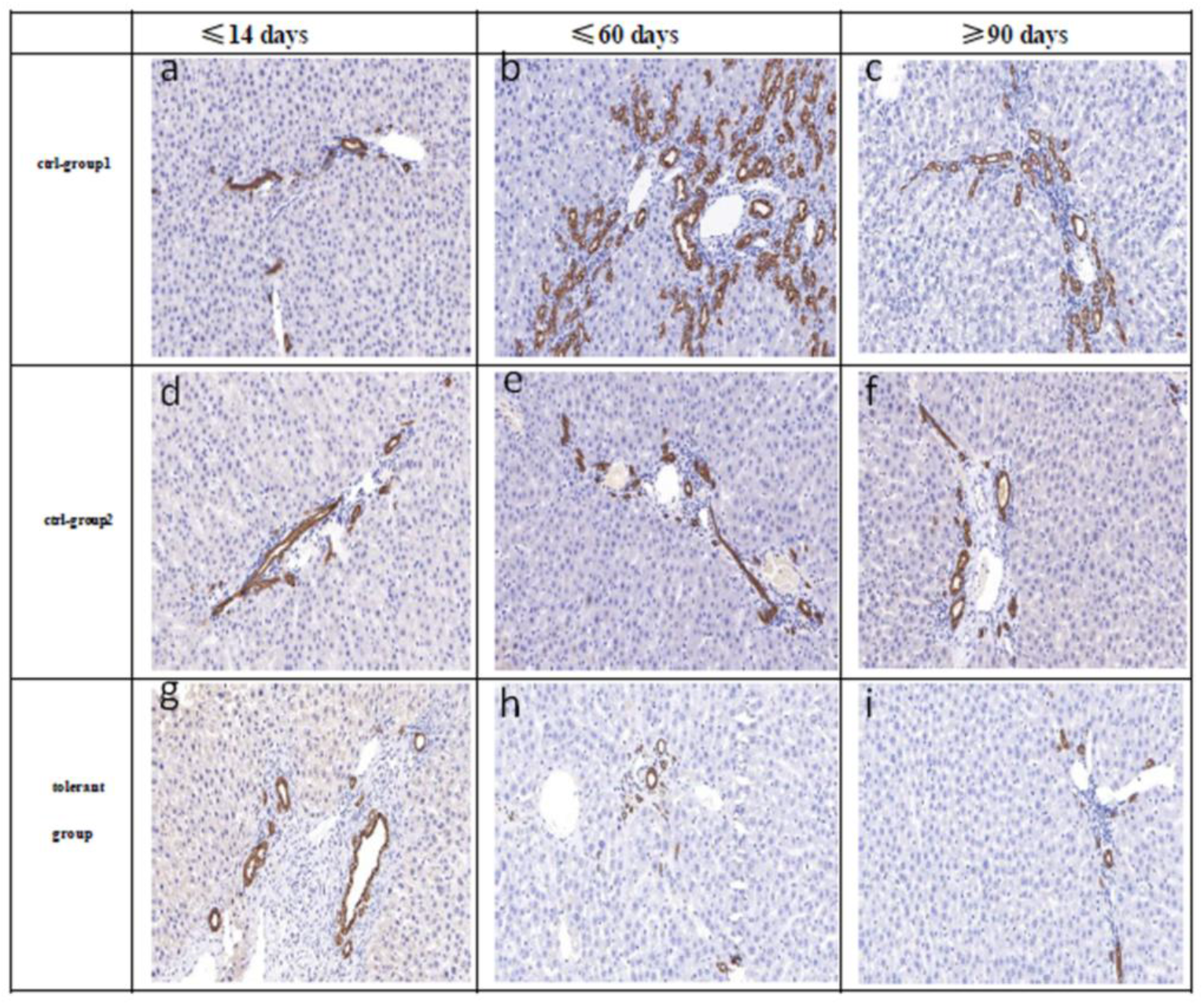
Histoimmunochemical staining for ductular reaction biomarker CK19. CK19 was most remarkable in the whole graft group(a-c,6,43,123 d), also remarkable in the half-size graft control group(d-f, 7,59,157d), not remarkable in the tolerant rats(g-I, 7,54,365d)

### Liver allograft remodeling during liver regeneration

To further gain insight into the underlying mechanism of liver regeneration in tolerance, we evaluated whether and how host-derived stem cells migrated, repopulated in the graft. 5-Bromo-2-deoxyuridine (Brdu) was used to evaluate liver regeneration, for the whole graft control group, liver regeneration almost discontinued; while it was relatively common in the tolerant grafts and the controlled half size allografts but much less frequent (Fig.6). CD34 as a biomarker of bone marrow stem cells was used to evaluate whether host-derived stem cells migrated and repopulated after r-GSF mobilization. Immunohistochemistry showed that CD34 was most pronounced in the allograft of tolerant rats (Fig.7), and that it was more pronounced in the control half size graft group than that of whole graft rats, indicating bone marrow stem cells repopulation was significant. CD133 was not detected in all allografts.

**Fig.6.**
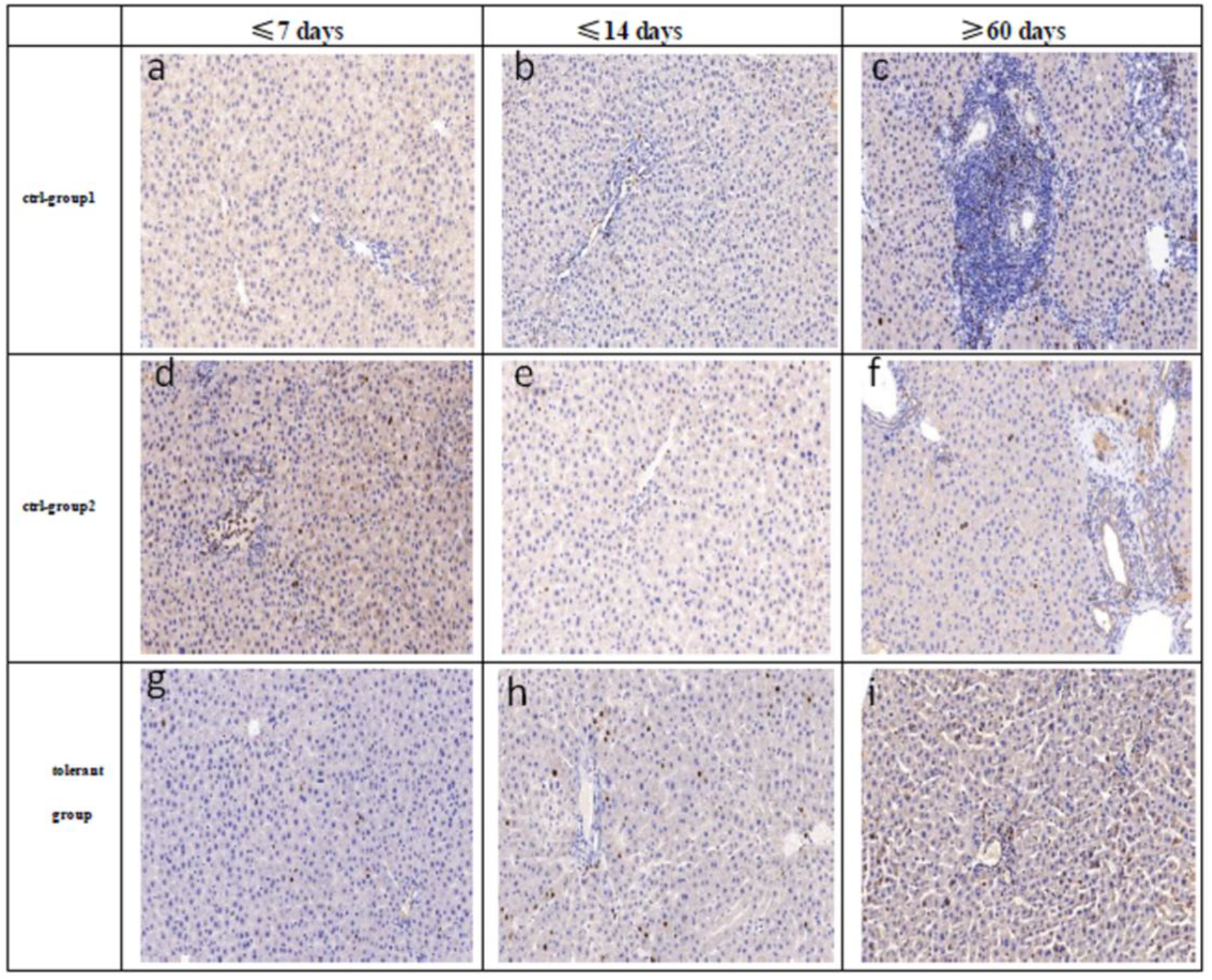
Liver regeneration in the allografts of 3 groups evaluated by BrDu in different periods. whole liver allograft showed almost no regeneration due to no injury (no signal), in the meantime, Brdu detection was more common in half-size allograft groups than that in the whole allograft(a-c,7,14,73d); after discontinued immunosuppression, immune injury developed insidiously in the whole liver allograft, on the contrary, Brdu detection was more common than that in the control half size rats (d-f, 6,14,157d) and the tolerant rats (g-i,7, 13, 365d).

**Fig.7.**
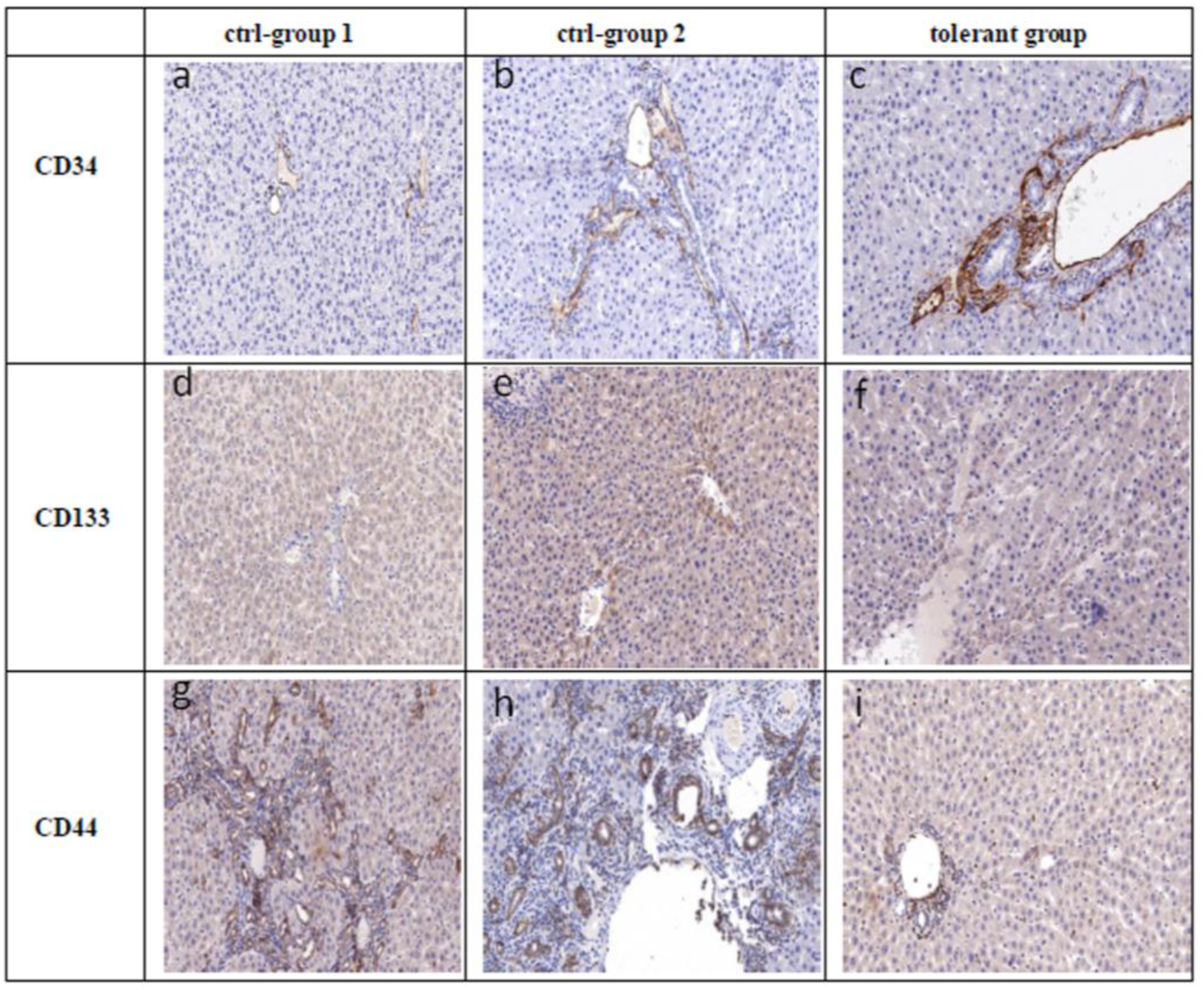
Peri-operatively histoimmunochemical staining for bone marrow stem cells with biomarkers of CD34, CD133. CD34 was most remarkable in the tolerant rats, more in the half-size graft of the control groups, less in the whole graft group. a(6 d),b(7d),c(7d).CD133 was negative in 3 groups, d(6d),e(7d),f(7d).CD44 stands for small hepatocytes. g (59 d),h (68d),I (341d).

### Donor-specific immune response

Lastly we perform skin transplantation to evaluate whether donor-specific immune hyporesponse occurs. Inbred donor strain skin transplantations were performed after 1 month post-LT. In animals of 3 groups, the skin allografts hardened and slough off in 2 weeks, robust immune reaction occurred to reject the skin allografts, indicating that naïve T cells failed to differentiate into memory T cells due to different spatial conformation of MHC molecules at the hypertrophy stage (Fig. 8).

**Fig.8.**
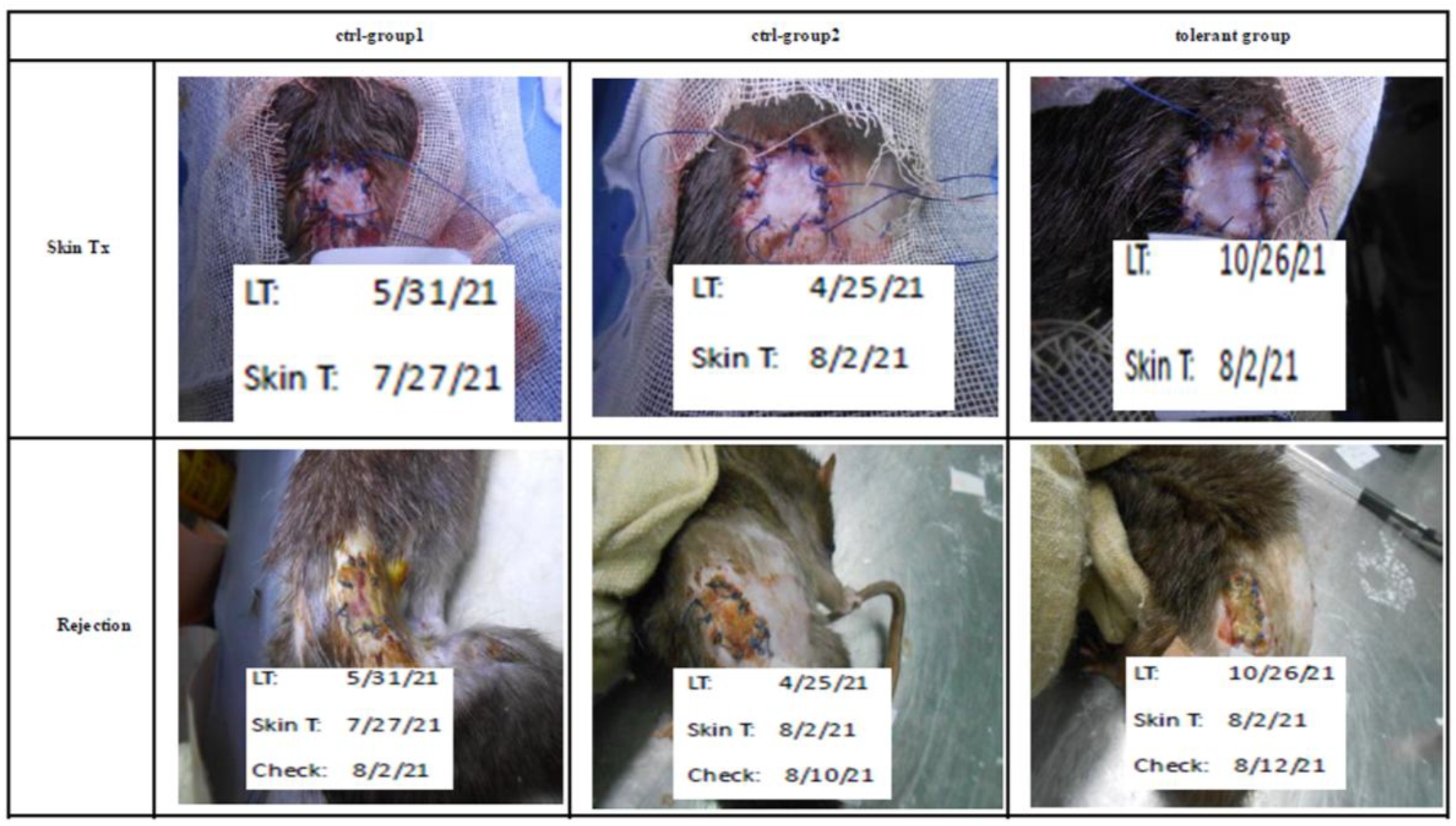
Skin transplantation for 3 groups. Donor skin grafts started to harden 4 days after operation and sloughed off in 14 days in 3 groups, indicating strong rejection occurrence.

## Discussion

After LT, long-lasting calcineurin inhibitor-based multiple regimen with narrow therapeutic window is required exclusively to protect the graft against the host immune destruction, in the upper of this window, their cumulative side-effects and toxicities are possibly associated with infection, de novo malignancy and metabolic syndrome^19,20^, it was reported that approximately 80% of LT recipients were over-immunosuppressed^21^; in the lower of this window, the allograft loss insidiously develops due to immune injury in the form of acute or chronic rejection. No optimized immunosuppression regimen is universally applicable. The side-effects and toxicities are the main roadblock of further improved outcomes of transplantation. Operational tolerance has emerged as the key strategy to further transplant outcomes.

Clinically spontaneous tolerance was reported for the recipients several years after LT, under one-fourth recipients remained off immunosuppression use, especially pediatric LT recipients whose partial or reduced size grafts were used^12,22^. It was suggestive that the unique liver regeneration phenomenon was possibly the mechanism of tolerance. Sun et al confirmed that plerixafor-mobilized bone marrow stem cells reinstituted the graft to prolong survival in half size graft LT model from Dark-Agouti (DA) to Lewis confirmed through Y chromosome track technique and the green fluorescent protein expression, it followed that CD133(not CD34) stem cells migrated and repopulated in the allograft. Their conclusion was self-perpetuating antigen specific immunosuppression rather than immune tolerance^23,24^. We speculated that half size DA liver graft underwent longer duration of regeneration because this rat is a poor metabolizer lacking cytochrome P450 subfamilies^25^, this model is the strongest rejection reaction resembling human LT, insufficient tacrolimus use (0.1mg/kg, totally four times in the first week) was the fundamental cause of chronic immune injury, resulting in compromised hepatocyte proliferation capacity, the small hepatocytes as one of hepatic progenitors started to proliferate, their study provided the clue as to that the injuries were ongoing in the allografts and liver regeneration or remodeling of the graft was taking place all the time.

Our protocol offers the operational tolerance confirmed with histological examination completely in accordance to Banff criteria^18^. Why is the smaller liver allograft harbored well by the host on the synergistic effect of liver regeneration and bone marrow stem cells? The truths need to be elucidated. We propose that a series of events after half size graft implantation play roles of tolerance achievement in our study. One immediate outcome after smaller allograft implantation is channeling the entire portal blood flow through half vessels to increase portal blood pressure and provide initiating signals for liver regeneration^26^. On ischemia reperfusion injury, hepatocytes would first undergo growth or hypertrophy as compensation and their volume would increase by half in a few days which was evidenced and supported by less Brdu detection in the half size allograft of our study. At the hypertrophy stage, the dynamic and subtle variation of spatial conformation of MHC molecules changed the interaction between MHC and antigen peptides and led to the failure of the recognition of MHC molecule antigen peptide complexes by T cell receptor’s exquisite docking ^27.28^. Bone marrow stem cells mobilized commonly with r-GSF will migrate, repopulate and differentiate in the half size graft; mesenchymal stem cells are immunosuppressive by downregulating expression of costimulatory molecules, inhibiting differentiation of dendritic cells from CD34 progenitors and reducing proinflammatory cytokine secretion^29^, as a result, tissue repair is improved quickly, and the allograft was tolerated. It seems to be more in line with the Danger model^30^. After hypertrophy switches to hyperplasia, spatial conformation of MHC molecule is not restored to their original because of the exquisite specificity. The immune recognition does not occur. Hepatocyte proliferative capacity restores, small hepatocyte’s specific biomarker CD44 were detected negative in tolerant rats, positive in control groups in our study^31^.

In our study, operational tolerance was achieved in 100% of hosts, the allograft bespoke the almost completely normal microstructures over one year, hepatic infiltration in the tolerant rats was consistent with outcomes reported by Taubert et al and Todo et al but less mild than their results^9,31^. Of note, our protocol is of highly clinical relevance that it targets the allograft and interventions are easily operable and reproducible: reduced-size or split LT are extensively performed in many centers; the host-derived bone marrow stem cells are attainably interventional; the short-course and low dose immunosuppressant use confer little impact on the hosts, and overarching side-effects and toxicities associated with immunosuppression can be waived. In view of liver regeneration to induce tolerance, its proliferative index can be used as biomarkers to predict the tolerant state. In addition, the graft can be divided into parts which will be supplied to 2 recipients; the paradigm of one liver for two recipients will surely expand the donor pool to shorten the waitlist of LT.

This operational tolerance was successfully induced in small animal, its procedures and intervention mimic human LT, nonhuman primate LT are warranted to expedite this protocol implicated in human LT.

## Abbreviations

LT: liver transplantation
MHC: major histocompatibility complex
ALT: Glutamic-pyruvic transaminase
AST: glutamic-oxaloacetic transaminase
Brdu: 5-Bromo-2-deoxyuridine

## Online content

Any method, additional references, Nature research reporting summaries, source data, extended data, supplementary information, acknowledgements are available online.

## Methods

### Data reporting

The sample size was predetermined according to the references without statistical methods^5,32,33^. The experiments were randomized and the investigators were blind during data analysis for all experiments.

### Rats

Rats (200-300g weight) including Sprague Dawley (SD), Lewis, Brown Norway (BN), served as donors and recipients purchased from Beijing Vital River Laboratory Animal Technology Corporation. All animals were housed and cared in a temperature and light-controlled environment, freely access to food and bottle water, they were fasted 12 hours before LT.

### Orthotopic rat LT and experimental grouping

Rat LT was performed by one surgeon. Under a modified mask for isoflurane inhalation anesthesia. Liver grafts were first perfused via the aorta with 5 ml of heparinized (50 U/mL) normal saline and then through the portal vein with 10 ml of cold lactated Ringer solution. The cold storage time was less than 3 hours. The recipient’s native liver was explanted out and the allograft was orthotopically implanted, followed by the supra-hepatic vena cava anastomosis with a 8-0 running suture. The two-cuff technique was used for the reconnection of the portal vein and the infra-hepatic vena cava^34,35^. The arterial reconnection was made with a stent^36^. The biliary continuity was achieved by a polyethylene stent. The recipients were monitored daily for mobility, posture, weight, fur, skin and urination color. 50% LT graft consists of removal of left lateral lobe, removal of left portion of the middle lobe and caudate lobes; for 70% graft, only left lateral lobe is removed^37^.

To make investigation of tolerance induction and the potential mechanism involved in liver regeneration, we performed OLT from Lewis to BN rat in the different groups, which is well-established model of strong acute rejection: whole graft LTs were performed to compare with half-size allograft LTs with daily subcutaneous injection of CSA at 2mg/kg for 9 days and r-GSF at 200u/kg for 5 days to study the significance of liver regeneration, half-size allograft LTs were performed only with same injection of CSA to emphasize the contribution of the host bone marrow stem cells into regenerating liver compared with tolerant group. 70% graft LTs were performed to compare with 50% graft LT for determination of graft size. Syngeneic graft LT was performed to provide the controls of histology images.

### Skin transplantation

Inbred Lewis rats were used as skin donors and LT BN rats used as recipients. A 1.5 cm × 1.5 cm full thickness graft was prepared from the donor abdomen; A graft bed (1.5 cm × 1.5 cm) was prepared on the back of a recipient rat^38^. When the skin graft was attached to the back of the recipient with interrupted sutures of 5-0 silk. The graft was covered with a protective tape and the first inspection was done 3 days after skin grafting and daily thereafter. Rejection was defined as skin graft acquired a red–brown color, hard consistency and slough.

### Evaluation of liver regeneration

5-bromo-2’-deoxyuridine (BrdU) was injected intraperitoneally at 50mg/kg 24 hour before liver tissue harvest. The liver tissues were then cut into small pieces that were fixed in 10% neutral formalin, embedded in paraffin, and sectioned. BrdU-incorporated hepatocytes were detected by use of an immunochemical system18 for monitoring cell proliferation with a monoclonal anti-Brdu cell proliferation kit^39^ (GB12051, Servicebio Inc. Wuhan, China).

### Blood collection and liver sample preparation

When the subject rat was euthanatized, the rat’s abdominal cavity was opened through the midline incision, the intestine was pulled out to expose the abdominal aorta, a blood collection needle was pierced into the aorta and connected to a negative air pressure tube containing EDTA for anti-coagulation, then the tube was upside down several times to prevent blood clotting. The plasma was gained at 2000g centrifuge for 10 minutes and aliquot was labeled and cryopreserved until use. According to spectrometry, the corresponding parameters were set on the automatic biochemical analyzer, the results were output after plasma is loaded.

The liver allograft was perfused with normal saline through the mesenteric vein and extracted out. The graft was weighed and cut into several small parts, then immerged in 10% neutral paraformaldehyde solution for later use.

### Hematoxylin-eosin and immunohistochemistry staining

Hematoxylin-eosin and masson staining were performed as conventionally described. Five-micrometer serially cut and frozen sections were fixed, the tissue section was placed in a retrieval box filled with citric acid (PH6.0) which was in a microwave oven for antigen retrieval to block endogenous peroxidase with 3% hydrogen peroxide solution. Then drop 3% BSA solution in the histochemical circle to cover the tissue evenly, and block for 30 minutes at room temperature. The primary antibodies were blocked with goat-derived rabbit serum, and other sources were blocked with BSA. The primary antibodies were added and incubated overnight. The secondary antibody (CD133) marked with HRP were added, whereas they were colored with DAB and controlled under microscope. Cell nucleus were counter-stained with hematoxylin. This procedures were applied to CD34,CD44,CK19 (Servicebio Inc. Wuhan, China)^24,40–43^.

### Statistics and reproducibility

Statistical analysis was performed using IBM SPSS Statistic 22.

### Data availability

All data generated and supporting the findings of this study are available within the paper, Additional information and materials will be made available including operation video upon request.

## Acknowledgements

This work was sponsored by Doctor initiative fund and project 23456 of Henan Provincial People’s Hospital.

## Authors’ contributions

GY C. funded and discussed the study. G.F.T. discussed the draft. H.B.Z. partly performed the experiments. S.D.W. performed the analysis of data. X.Y.G animal caring. F.Z.L. discussed the project. D.L. collected data. H.G. designed and performed the pathological examinations. S.T. Z. conceived, designed, carried out, funded the experiment and performed OLT, wrote manuscript and finalized the study.

---

**Extended Data Table.1.** Experimental grouping for different graft size, treatment and survival. Cyclosporine A: CSA; BMSC: bone marrow stem cells, this was to provide the references and comparison of histology images.

| Combinations | Graft | N | Treatment | Survival (days) |
| --- | --- | --- | --- | --- |
| Lewis→Lewis | 50% | 2 | no | 156, 214 |
| SD → Lewis | 70% | 3 | CSA+BMSC | 185,183,178 |
| Lewis→BN | 70% | 2 | CSA+BMSC | 124,152 |

**Extended Data Fig.1.**
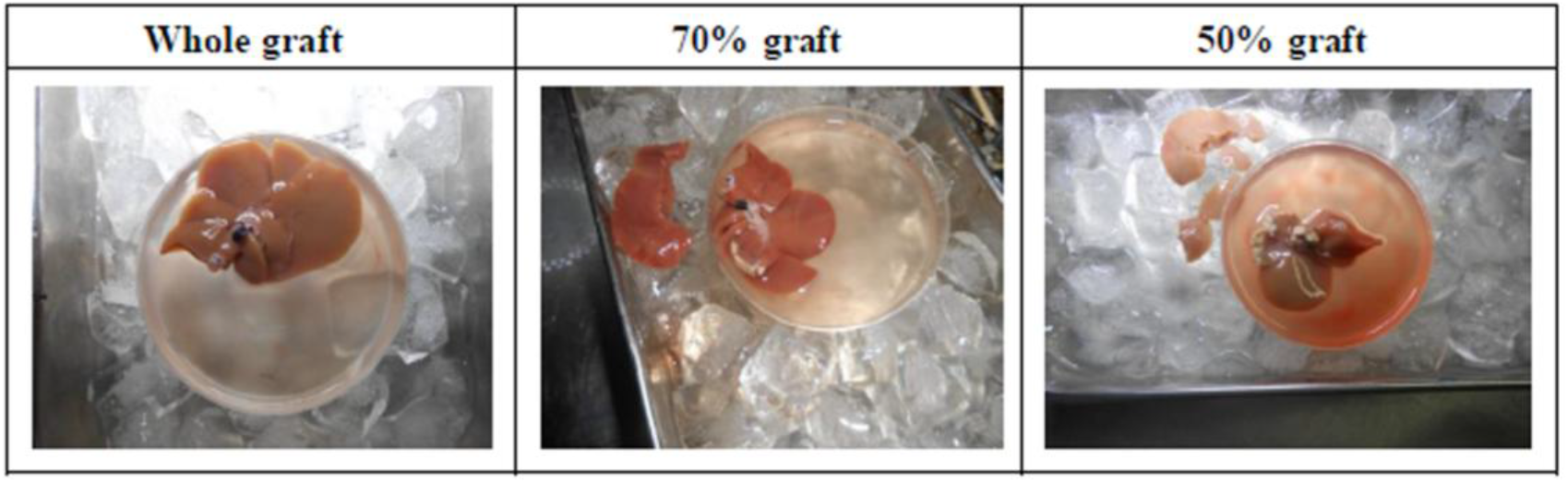
The preparation of graft size in different groups. The removed lobes were shown.

**Extended Data Fig. 2.**
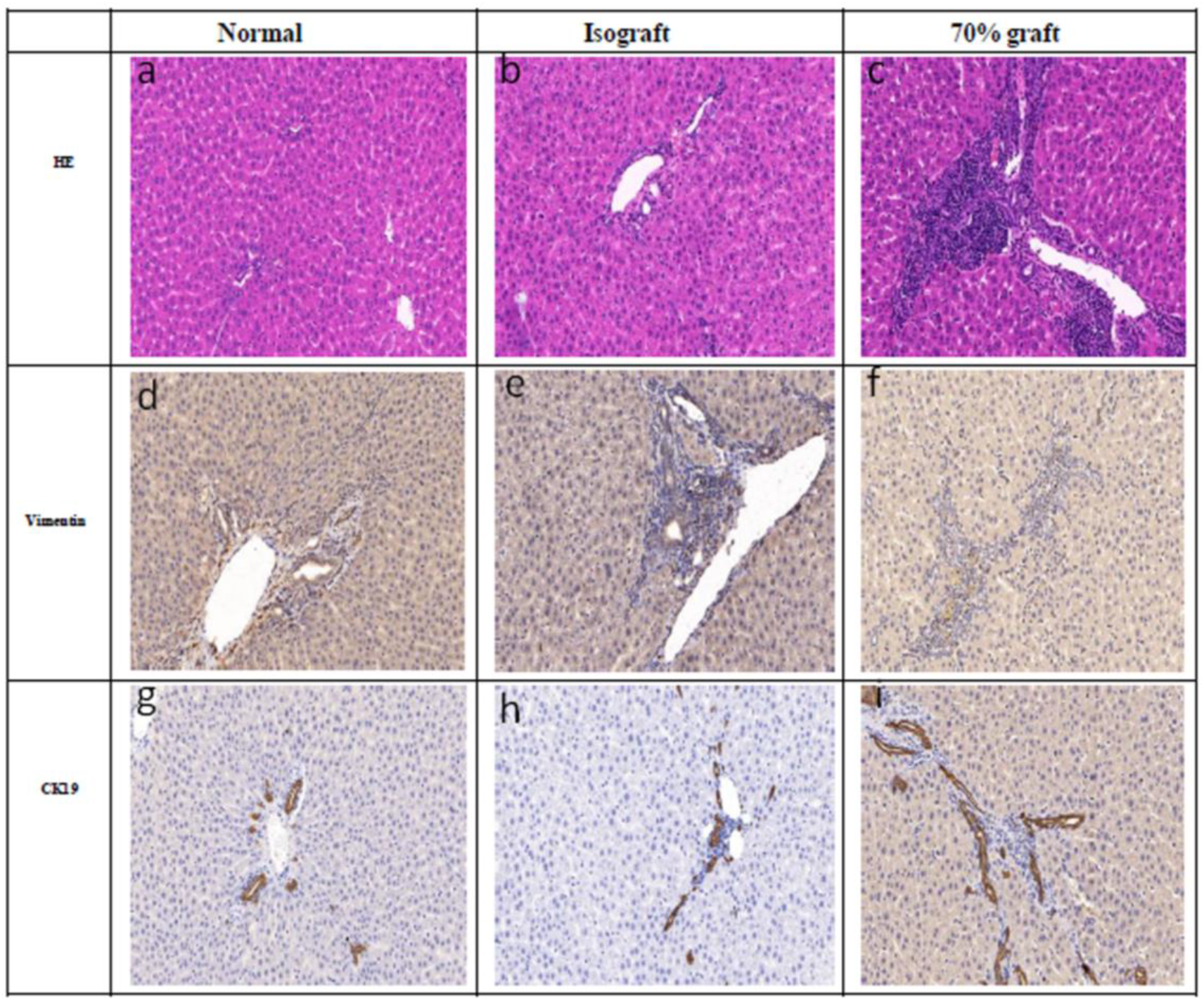
Histology images of the normal rats, the syngeneic graft rats and 70% graft used for reference. Normal rats were from the host native livers. a-c for HE of different groups, b (214d),c (183d). d-f for vimentin, e (156d), f (183d). g-i for CK19, h (156d), I (183d).

